# Context-dependent hyperactivity in *syngap1a* and *syngap1b* zebrafish autism models

**DOI:** 10.1101/2023.09.20.557316

**Authors:** Sureni H. Sumathipala, Suha Khan, Robert A. Kozol, Yoichi Araki, Sheyum Syed, Richard L. Huganir, Julia E. Dallman

**Author notes:** Corresponding Author: Julia E. Dallman, Associate Professor of Biology, University of Miami.

## Abstract

**Background and Aims:** SYNGAP1 disorder is a prevalent genetic form of Autism Spectrum Disorder and Intellectual Disability (ASD/ID) and is caused by *de novo* or inherited mutations in one copy of the *SYNGAP1* gene. In addition to ASD/ID, SYNGAP1 disorder is associated with comorbid symptoms including treatment-resistant-epilepsy, sleep disturbances, and gastrointestinal distress. Mechanistic links between these diverse symptoms and *SYNGAP1* variants remain obscure, therefore, our goal was to generate a zebrafish model in which this range of symptoms can be studied.

**Methods:** We used CRISPR/Cas9 to introduce frameshift mutations in the *syngap1a* and *syngap1b* zebrafish duplicates (*syngap1ab*) and validated these stable models for Syngap1 loss-of-function. Because *SYNGAP1* is extensively spliced, we mapped splice variants to the two zebrafish *syngap1a* and *b* genes and identified mammalian-like isoforms. We then quantified locomotory behaviors in zebrafish syngap1ab larvae under three conditions that normally evoke different arousal states in wild type larvae: aversive, high-arousal acoustic, medium-arousal dark, and low-arousal light stimuli.

**Results:** We show that CRISPR/Cas9 indels in zebrafish *syngap1a* and *syngap1b* produced loss-of-function alleles at RNA and protein levels. Our analyses of zebrafish Syngap1 isoforms showed that, as in mammals, zebrafish Syngap1 N- and C-termini are extensively spliced. We identified a zebrafish *syngap1* α1-like variant that maps exclusively to the *syngap1b* gene. Quantifying locomotor behaviors showed that *syngap1ab* larvae are hyperactive compared to wild type but to differing degrees depending on the stimulus. Hyperactivity was most pronounced in low arousal settings, with overall movement increasing with the number of mutant *syngap1* alleles.

**Conclusions:** Our data support mutations in zebrafish *syngap1ab* as causal for hyperactivity associated with elevated arousal that is especially pronounced in low-arousal environments.

## Background

SYNGAP1 syndrome is caused by genetic variants in the *SYNGAP1* gene and is one of the most prevalent genetic forms of intellectual disability (ID) (Hamdan et al., 2010; Hamdan et al., 2011; Berryer et al., 2013; Satterstrom et al., 2020). In addition to ID, people with SYNGAP1 syndrome present with Autism Spectrum Disorder, epilepsy, gastrointestinal distress, developmental delay, hypersensitivity to sound and light, high pain thresholds, and increased risk-taking (Pinto et al., 2010; Carvill et al., 2013; Kilinc et al., 2018; Weldon et al., 2018; Jimenez-Gomez et al., 2019; Vlaskamp et al., 2019). The majority of SYNGAP1 syndrome-causing variants are *de novo*, occurring in the child but not in their parents (Hamdan et al., 2010; Hamdan et al., 2011; Berryer et al., 2013); SYNGAP1 syndrome is caused by haploinsufficiency, therefore, having one *SYNGAP1* variant is sufficient to cause symptoms.

To better understand genotype/phenotype relationships in SYNGAP1 syndrome, we generated loss-of-function mutations in zebrafish *syngap1a* and *syngap1b* duplicates using CRISPR/cas9 (Varshney et al., 2016). With accessible early development, optically transparent embryos, high fecundity, and established methods for genetic manipulation, zebrafish complement extant rodent models to understand the role of disease genes in development and behavior (Kozol et al., 2016; Sakai et al., 2018). While mammals have a single *SYNGAP1*, zebrafish *syngap1* is duplicated due to a whole genome duplication event 50-80 million years ago and retention of both *syngap1a* and *syngap1b* (Glasauer and Neuhauss, 2014; Kozol et al., 2016). Two recent papers generated zebrafish *syngap1b* models that differ from the model we report here because they only target one of the duplicate zebrafish *syngap1* genes (Colon-Rodriguez et al., 2020; Griffin et al., 2021).

Mammalian *SYNGAP1* mRNAs are extensively spliced at N- and C- termini and the C-terminal isoforms α1, α2, β, and γ have been functionally characterized (McMahon et al., 2012; Araki et al., 2020; Gou et al., 2020; Kilinc et al., 2022). The α1 isoform is highly enriched at the post-synaptic density of glutamatergic synapses through a four amino-acid PDZ-interacting domain by which it interacts with the synaptic scaffolding protein PSD-95 (Chen et al., 1998; Kim et al., 1998; Komiyama et al., 2002). Here we annotate zebrafish mRNAs for how they might correspond to these mammalian isoforms and map zebrafish isoforms to *syngap1a* and *syngap1b* genes.

Individuals with SYNGAP1 syndrome show sensory hyperactivity, sometimes even seizures, in response to sensory stimuli such as eating, light, sound, touch and/or pain. These symptoms are a major concern of the SYNGAP1 parents and caregivers because they can put these individual’s lives at risk (Vlaskamp and Scheffer, 2020; Lyons-Warren et al., 2022). Consistent with the human symptoms, rodent models also show sensory-induced hyperactivity as well as seizures that can be induced by loud sounds (Guo et al., 2009; Ozkan et al., 2014; Michaelson et al., 2018; Creson et al., 2019; Sullivan et al., 2020). To assess sensory-induced behaviors in *syngap1ab* zebrafish mutants, we used two standard sensorimotor assays: vibration to evoke the acoustic startle response (ASR) and transitions between light and dark to evoke the visual-motor response (VMR) (Emran et al., 2008; Gao et al., 2014; Dunn et al., 2016). Because changes to sensory habituation could contribute to sensory processing issues in SYNGAP1 patients (Tavassoli et al., 2014; Robertson and Baron-Cohen, 2017; Oldehinkel et al., 2019), we also conducted a short-term habituation assay to see how the Syngap1ab zebrafish larvae behave towards supra-threshold stimuli that are presented in rapid succession.

Like mammalian models, our zebrafish *syngap1ab* mutant models exhibit hyperactivity in both ASR and VMR assays. Hyperactivity was least pronounced in response to aversive acoustic stimuli and most pronounced during low-arousal light conditions. By analyzing the frequency distributions of movement distance and rest duration, we show that *syngap1ab* model hyperactivity is due to higher-frequency, larger movements that resemble goal-directed behaviors associated with heightened states of arousal.

## METHODS

### Fish Maintenance and Husbandry

All the zebrafish larvae and adults used in this study were reared at the University of Miami Zebrafish facility as per IACUC protocol #18-129. Water temperatures were maintained at 28 °C and both adult and larval zebrafish were exposed to a circadian cycle of 14 hours light/10 hours dark. Water housing adult zebrafish was continuously monitored for pH and conductivity to maintain optimal conditions. Upon collection, embryos were reared in 10 cm petri dishes that were cleaned daily to prevent fungal growth that could limit oxygen and stunt early embryonic growth. Larvae used for behavioral assays were raised ∼50 larvae per petri dish to minimize competition and developmental delays.

### CRISPR/Cas9 generation of syngap1ab mutant zebrafish

*syngap1ab* mutants were generated using CRISPR/Cas9 genome editing technology. One cell stage WT embryos were injected with small guide RNA (sgRNA; Integrated DNA Technologies-IDT) designed to target exon 4 of *syngap1a* gene and exon 5 of *syngap1b* gene, along with Cas9 protein in a 1:5 ratio. To minimize sgRNA competition and for sequencing purposes, each embryo was injected with either *syngap1a* or *syngap1b* sgRNA. Resulting mosaic F0 larvae were reared to adulthood and crossed to wild type animals to generate an F1 generation for sequencing to identify *syngap1a* and *syngap1b* mutant alleles with indels resulting from CRISPR editing (see below). Upon identification of the mutant *syngap1a* and *syngap1b* alleles, they were in-crossed to obtain *syngap1a*b double mutants. Adult *syngap1a*b mutant fish span F2-F5 generations with various *syngap1a* and *syngap1b* allele combinations. For the following experiments, adult male *syngap1ab-/-* were outcrossed to WT (AB/TL https://zfin.org/action/genotype/view/ZDB-GENO-031202-1) females to obtain *syngap1a*b+/- larvae and were in-crossed with *syngap1ab-/-* to obtain *syngap1ab-/-* larvae. For simplicity, zebrafish that are heterozygous in both *syngap1a* and *syngap1b* genes are denoted as *syngap1ab+/-*, and those homozygous in both *syngap1a* and *syngap1b* genes are denoted as *syngap1ab-/-.* All the wild type (WT) larvae used for this study were AB/TL unless otherwise stated and are denoted as *syngap1ab+/+*.

### syngap1ab alleles used in this study

The molecular identity of CRISPR alleles was determined by obtaining a small caudal fin sample from each anesthetized (200 mg/ L of Tricaine (MS-222)) F1 adult fish. Genomic DNA was extracted by digesting each fin sample in 50 µl of 50 mM NaOH at 95 °C for 1 hour. When larval genotypes were to be determined, each larva was anesthetized by either placing them on ice for 30 mins or by using 200 mg/L Tricaine (MS-222). Upon complete anesthetization, larval genomic DNA (gDNA) samples were isolated using the previously described method.

Gene specific primers (Table 1) for both genes were designed using Primer3 software. Primer3 input sequence for each gene was selected based on their Cas9 target regions. For PCR, each reaction mixture contained 5µl of 10x GOTaq Polymerase (Promega), 0.5 µl from each 10 µM *syngap1a* forward primer and reverse primers or 0.5 µl from each 10 µM *syngap1b* forward primer and reverse primers (IDT), 3 µl of nuclease-free water, and 1 µl of gDNA (from either larvae or fin clip digestions). Resulting PCR products were sent out for sequencing (Eurofins Genomics, LLC) and the results were read and analyzed using the ApE – A plasmid Editor v2.0.61 (Davis and Jorgensen, 2022) and SnapGene viewer software to determine the *syngap1a* and *syngap1b* mutant alleles (Supplementary Figure 1).

**Table 1.**
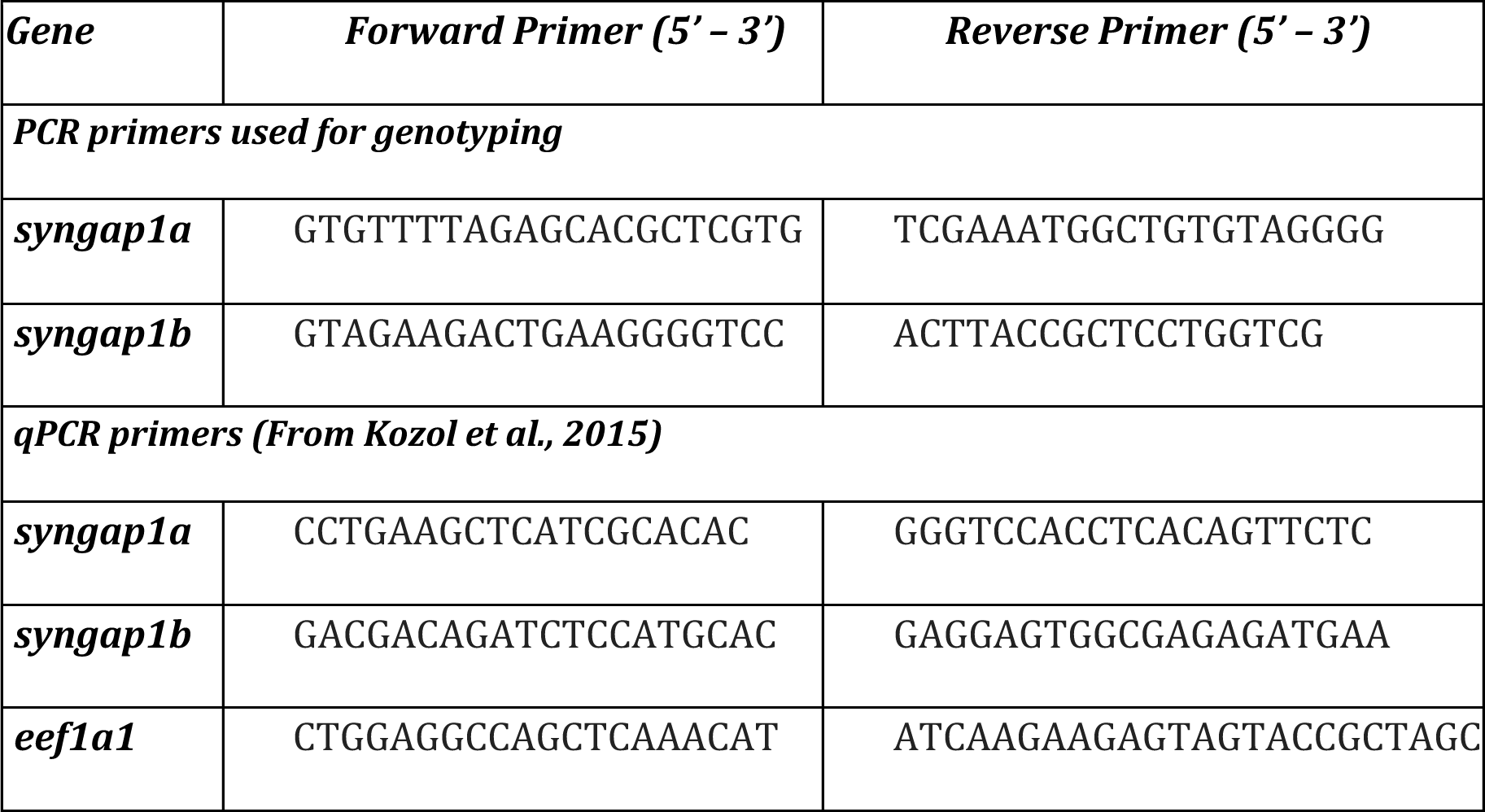
*syngap1* primers used for genotyping and qPCR.

### syngap1a and syngap1b isoform identification

To determine the zebrafish *syngap1a*b splice variants/ isoforms, mRNA sequences (both published and predicted), were obtained from the NCBI protein database. To test for evidence of isoform expression, these sequences were searched against Expressed Sequence Tags (EST) and Transcriptome Shotgun Assembly (TSA) databases. Expressed isoforms were then BLASTed against the UCSC zebrafish genome browser to identify unique, common, and alternative splice variants to better annotate available *syngap1a*b isoforms (Fig. 1).

**Figure 1.**
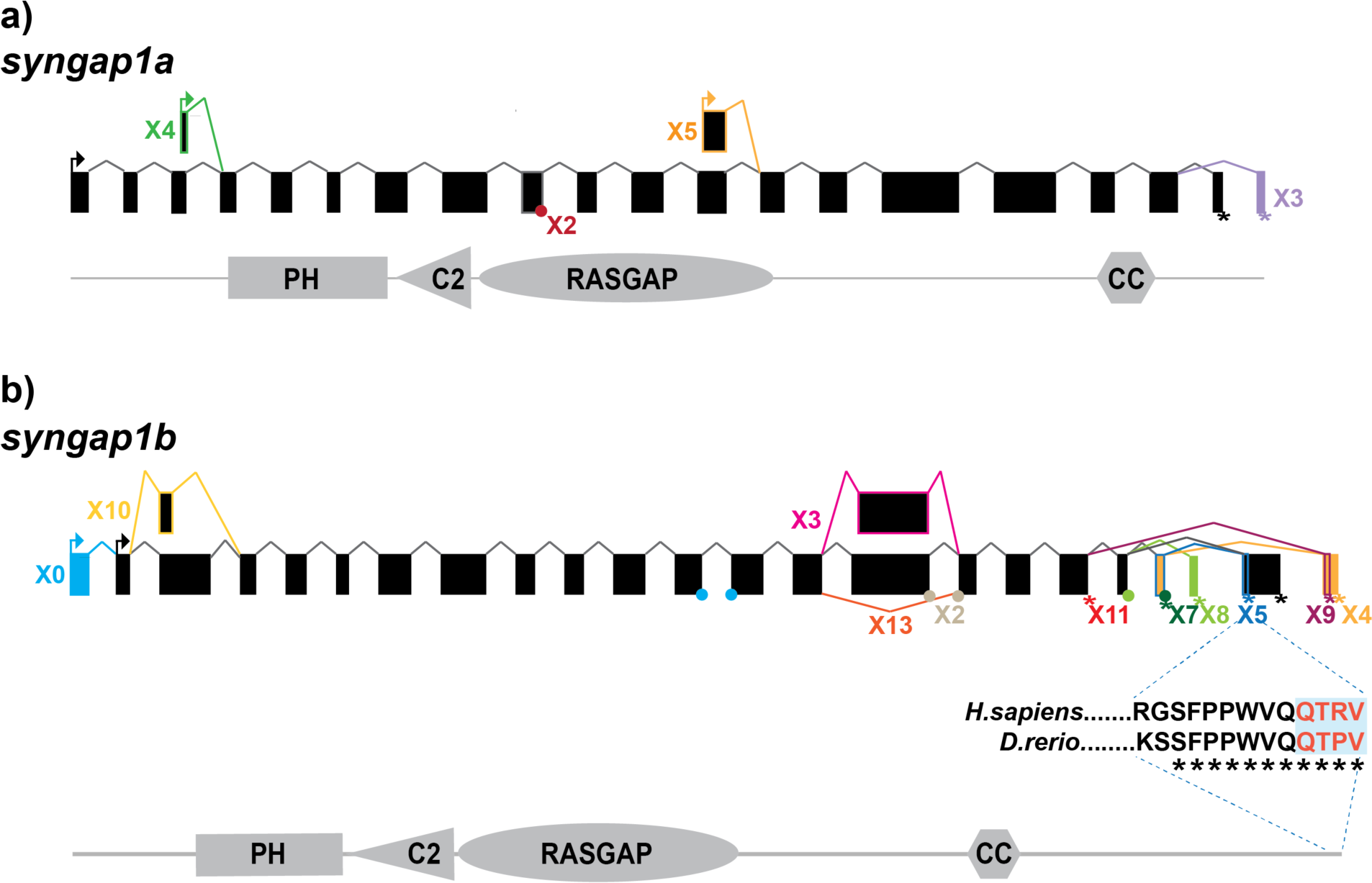
Zebrafish *syngap1a* and *syngap1b* isoforms. NCBI database search revealed **a)** five *syngap1a* (X1 - X5) and **b)** eleven *syngap1b* isoforms (X0, X2-5, X7-11, & X13). Alternative splice sites are shown either using a dot (if the exon is missing <10 bp) or a box (if the difference is >10 bp). Similar box colors with colored connecting lines suggest that they share similar nucleotide sequences but will have cryptic splice sites that the splicing mechanism would recognize. Stop codons are shown using an asterisk (*) with their respective colors.

### qPCR

To check for nonsense-mediated mRNA decay in *syngap1a*b mutants, qPCR was used to quantify relative *syngap1a*b expression levels in *syngap1a*b mutant larvae compared to WT larvae. To extract RNA, 7 days post-fertilization (dpf) old larvae were anesthetized by placing them on ice for 30 minutes. For RNA extractions, we used TRIzol (Life Technologies, Carlsbad, CA, USA). For each genotype (WT, *syngap1a*b+/-, and *syngap1a*b-/-), experiments were conducted in triplicate with twenty larvae pooled together per sample and 1 µg of RNA was used as input for RT-PCR. To enhance syngap1 cDNA in each sample, *syngap1a* and *syngap1b* gene specific primers were used with Eukaryotic translation elongation factor 1 like 1 *eef1a1l1* (ZFIN) as the internal control. cDNA was made using SuperScript III (Invitrogen™) and incubating at 50°C for 1 hour followed by 15 minutes at 70°C. qPCR was carried out using GoTaq qPCR Probe Kit (Promega, Cat #A6110) in a QuantStudio3 RT-PCR system (Applied Biosystems™) based on manufacturer’s protocols. Cycling conditions were as follows: Activation step of 95°C for 10 minutes, followed by PCR with 40 cycles of 95°C for 15 seconds and 60°C for 1 minute, followed by a melt curve of 95°C for 15 seconds and 60°C for 1 minute. Relative levels of gene expression were calculated using the Δ1Δ1Ct method. For this method cycle threshold Ct values for *syngap1a* and *syngap1b* genes were first normalized to Ct values of the internal control *eef1a1l1* by calculating Δ1Ct: Ct_syngap1a_-Ct_eef1a1l1_ and Ct_syngap1b_-Ct_eef1a1l1_. Fold-changes in *syngap1* gene expression were then compared in WT, *syngap1ab+/-*, and *syngap1ab-/-* larvae by calculating 2^-ΔΔCt^ with ΔΔCt: ΔCt*_syngap1ab_* _mutant_-ΔCt_WT_. Fold changes in *syngap1ab* mutants were calculated by dividing WT values.

### Western blot analysis

Brain regions (mouse) or whole brains (zebrafish) were excised from C57BL6 mice or adult zebrafish. Tissues were lysed in 10 volumes (for zebrafish, each brain was considered ∼15 mg) of lysis buffer (50 mM Tris pH 8.0, 100 mM NaCl, 1 mM EDTA, 1 mM EGTA, 1% TritonX-100, 0.2% SDS, 0.5% Sodium deoxycholate, with complete Protease inhibitor EDTA-free mix (Roche/SIGMA)) using a Dounce homogenizer to obtain homogenized tissue samples. Each sample was then diluted by 1:10 using lysis buffer and ∼20 µl from each sample was loaded into each gel lane. Based on these experimental settings, each sample i.e. for both zebrafish and mouse tissue samples, contained about 29 µg of proteins per 20 µl. Samples were first probed with SYNGAP1 antibodies (Abcam ab3344 or NOVUS nbp2-27541) and were followed by probing with secondary antibodies (anti-rabbit IgG IRDye680; LICOR 926-68071 or anti-goat IgG IRDye680; LICOR 926-68074). Resulting signals were measured and imaged using a fluorescence-based imaging system (Odyssey CLx Imaging System).

### Visual Motor Response (VMR) assay

6 dpf larvae were used for all the behavioral assays. *syngap1ab* mutants and WT larvae were placed in a 96-square well plate containing system water. Prior to starting the experiment, larvae were dark adapted for 1 hour and were maintained at 28 °C in the Noldus DanioVision^TM^ behavioral observation chamber. Ethovision® XT 11 software was used to program the delivery of stimuli and to analyze the results. Images were captured at 40 Hz. For the visual-motor response (VMR) assay, larvae were exposed to 5 minutes of 12% (high-light settings) light on stimulus followed by 5 minutes of light off stimulus. Larval movements were recorded for four consecutive light on/ off cycles and their activity/movements were recorded for a total duration of 40 minutes. Larvae were then genotyped using the genotyping assays previously described and analyzed for the total distance moved as a proxy for their larval activity for both lights on and off conditions. The resulting data was further analyzed using GraphPad Prism 9.1 and MATLAB software.

### Displacement and dwell-time analyses

To test whether the observed hyperactivity is due to increased initiation of movements or increased distance moved, we analyzed the displacement and dwell-time data from raw, exported data produced by Ethovision® XT 11 software and these were analyzed using MATLAB scripts (https://github.com/sheyums/Sureni_Sumathipala_syngap1.git) to examine behavioral frequency distributions. Resulting data were plotted using Prism GraphPad (V9.1).

### Short-term habituation assay

To test whether the larvae show habituation in response to acoustic stimuli, a short-term habituation assay was used as described in Wolman et al., 2015. After 30 minutes of light adaptation, 6 dpf WT and *syngap1ab+/-* larvae were presented with 5 phases of acoustic stimulation using a Noldus Daniovision observation chamber. Phase 1 consisted of 10 tap stimuli (intensity level 3) delivered at a 20 sec inter stimulus interval (ISI). Phase 2 consisted of 10 tap stimuli (intensity level 5) delivered at also at a 20 sec ISI. Phase 3, the habituation test, consisted of 30 tap stimuli (intensity level 5) delivered at 1 sec ISI. Phase 4 was a 3 minute rest period. Lastly phase 5 consisted of 10 tap stimuli (intensity level 5), delivered at a 20 sec ISI. Total distance moved by each larva per second was analyzed to measure short-term habituation. Data were analyzed using Microsoft Excel and Prism GraphPad software (V9.1).

## Results

### Zebrafish syngap1b but not syngap1a encodes an isoform that is similar to the mammalian PDZ-interacting Syngap1α1

While mammals including humans have a single *SYNGAP1* gene that encodes a 1343 amino acid, ∼149kDa protein (Komiyama et al., 2002), there are two *syngap1* genes present in the zebrafish genome. The zebrafish *syngap1a* gene is found on chromosome 16 and encodes a 1290 amino acid, ∼146kDa protein whereas the *syngap1b* gene is found on chromosome 19 and encodes a 1507 amino acid, ∼160kDa protein. Like rodents and humans, both zebrafish Syngap1a and Syngap1b have four, highly conserved protein-protein interacting domains: pleckstrin homology (PH), C2, RasGAP, and coiled-coiled (CC).

We characterized zebrafish Syngap1 isoforms based on NCBI databases and comparisons to mammalian isoforms. In mammalian Syngap1, there are four well-characterized, alternatively-spliced Syngap1 isoforms, α1, α2, β, and γ, that vary at their C-termini (McMahon et al., 2012). To assess whether zebrafish *syngap1a* and *syngap1b* genes might encode similar isoforms, we curated all published and predicted *syngap1ab* isoforms. Similar to the mammalian isoforms, for both zebrafish *syngap1a* and *syngap1b* genes, exons encoding the four protein interaction domains occur in all isoforms with many alternative exons at both N- and C-termini. Based on expressed sequence databases, we were able to identify five *syngap1a* and eleven *syngap1b* isoforms (Fig. 1). Of the human C-term isoforms, α1 is the most studied isoform and is localized to the post-synapse by a four amino acid (QTRV) PDZ-interacting domain. Interestingly, we were able to find a zebrafish *syngap1b* C-term isoform X5 with a putative mammalian-like PDZ-interacting domain that had eleven of twelve of the terminal amino acids identical to those of human (Fig. 1b). We were unable to find mammalian-like isoforms (α2, β, and γ) due to the highly variable C-terminal ends in the zebrafish isoforms.

### N-terminal CRISPR/Cas9-generated alleles produce loss-of-function SYNGAP1 models

To better understand how pathogenic mutations in *syngap1* result in altered behaviors, we used CRISPR/Cas9 to generate zebrafish loss-of-function mutations. In humans, *SYNGAP1* pathogenic variants span throughout the gene (Hamdan et al., 2011; Berryer et al., 2013; Vlaskamp et al., 2019; Gamache et al., 2020). To mutate a region of the gene that would affect all isoforms, we targeted the earliest shared exons, exon 4 and exon 5 in *syngap1a* and *syngap1b* respectively, to generate loss-of-function alleles. CRISPR/Cas9 induces indels causing reading frame shifts and introducing premature stop codons. Upon sequence analyses of F1 adult crispants, we selected two mutant alleles for *syngap1a* (*syngap1a* +7 nucleotide insertion and *syngap1a* -22 nucleotide deletion) and one allele for *syngap1b* with a -14 nucleotide deletion, all of which would be predicted to result in a severely truncated proteins that were less than 200aa (Fig. 2b). To best recapitulate human SYNGAP1 variant haploinsufficiency, we used double-heterozygous larvae for *syngap1a* +7 and *syngap1b* -14, and to further assess complete loss-of-function, we used double-homozygous larvae (here onwards denoted as *syngap1ab*+/- and *syngap1ab*-/- respectively).

**Figure 2.**
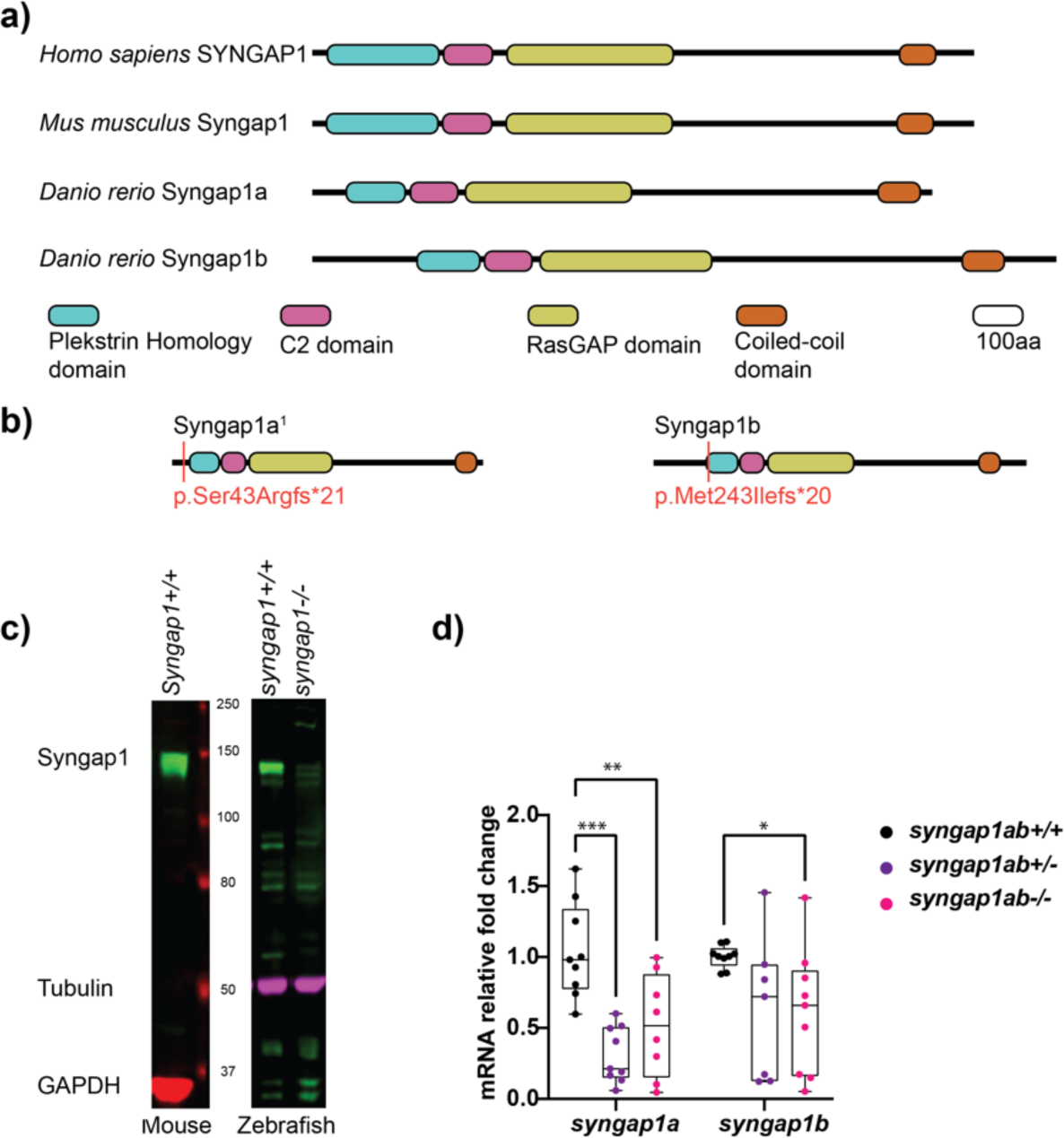
Zebrafish loss-of-function model for human SYNGAP1 syndrome. **a)** Mammalian SYNGAP1 protein (*H. sapiens* and *M. musculus*) has four main protein interacting domains; pleckstrin homology (PH) domain, C2 domain, RasGAP domain, and coiled-coiled(CC) domain. Zebrafish (*D. rerio*) Syngap1 orthologs; Syngap1a and Syngap1b also show domain conservation with that of mammals. **b)** Syngap1ab protein diagrams show sites of CRISPR induced mutations. Resulting CRISPR mutants used for phenotypic analyses: Syngap1a^1^ allele p.Ser43Argfs*21 and Syngap1b allele p.Met149Ilefs*9. Syngap1a^1^ had an amino acid change from a serine to an arginine at position 43 introducing a premature stop codon, 21 amino acids downstream. Syngap1b mutant allele had a change of methionine to an isoleucine at position 49 introducing a premature stop codon 9 amino acids downstream. **c)** Western Blots show SYNGAP1 expression in adult mouse and zebrafish whole brain lysates. WT zebrafish Syngap1 was identified at a similar size (∼150kDa) with that of mouse SYNGAP1 using rat anti-Syngap1. GAPDH and tubulin were used as the loading controls for mouse and zebrafish respectively. **d)** Mutant *syngap1ab* larvae showed reduced *syngap1* and *syngap1b* mRNA expression levels at 7dpf. Group comparisons were made using 2-way ANOVA followed by Tukey’s multiple comparison test. P value asterisks represent; p<0.05 - *, p<0.01 - **, p<0.001 - ***, p<0.0001-****

We tested for loss-of-function of *syngap1* at the level of protein and mRNA. Western blot analysis carried out using adult zebrafish brain lysates showed reduced Syngap1 protein levels in *syngap1a*b-/- brain tissues (Fig. 2c). qPCR analysis of RNA harvested from 7-day-old larvae showed reduced RNA transcripts in both *syngap1ab+/-* (adjusted p values for *syngap1a*= 0.001, and *syngap1b*=0.0495) and *syngap1ab-/-* (adjusted p values for *syngap1a*= 0.0699) and *syngap1b*=0.0312) mutant larvae compared to WT larvae supporting non-sense mediated decay (Fig. 2d). Taken together, these results show that our CRISPR/cas9 generated *Syngap1a*b zebrafish mutants show reduced mRNA and protein expression, supporting the use of these alleles as haploinsufficient models.

### Evidence for cooperativity between syngap1a and syngap1b genes for survival

To study how each syngap1 duplicate impacts survival and behavior, we took into consideration the overlapping mRNA expression levels of *syngap1a* and *syngap1b* during early development (Kozol et al., 2015) and analyzed larval survival from each of the nine genotypes resulting from *syngap1ab*+/- in-crosses. When *syngap1b* mutant alleles outnumbered *syngap1a* mutant alleles, larval survival was lower than expected (Supplementary Figure 2; p<0.0001). These observations suggest that there is a relationship between the *syngap1a*:*syngap1b* expression ratio and that the *syngap1b* ortholog is more important than *syngap1a* ortholog for larval survival when *syngap1b* mutant alleles outnumber those of *syngap1a*.

### Syngap1ab+/- larvae show greater dynamic range, elevated response probability, and normal habituation in response to acoustic stimuli

Given that SYNGAP1 is known to be important for sensory processing, we wanted to test the dynamic range of *syngap1ab+/-* zebrafish responses to low frequency vibration stimuli of medium and high intensity as well as their ability to habituate to repeated, high-intensity, high-frequency stimuli. To assess these factors, we used an assay described in Wolman et al., 2015 in which 6 days-post-fertilization (dpf) larvae were exposed to different intensity vibrations with different inter-stimulus intervals (ISIs) during phases 1-5 of the assay (Fig.3a). We allowed larvae to acclimate to the Noldus chamber for 30 minutes before exposing them to vibrational stimuli. During phases 1 and 2, 10 taps of medium-intensity (level 3) and high-intensity (level 5) respectively were delivered with 20 second ISI. During phase 3, habituation is tested by delivering 30 high intensity taps with one second ISI. This was followed by a three minute rest period in phase 4. Finally, phase 5 was a repeat of phase 2 with 10 high-intensity taps delivered with 20 second ISI. Overall, *syngap1+/-* larvae (n=130 from 3 independent crosses) responded very similarly to WT larvae (n=110 from three independent crosses) (Fig. 3b), with normal a degree of habituation to high frequency stimuli (Fig. 3c).

**Figure 3.**
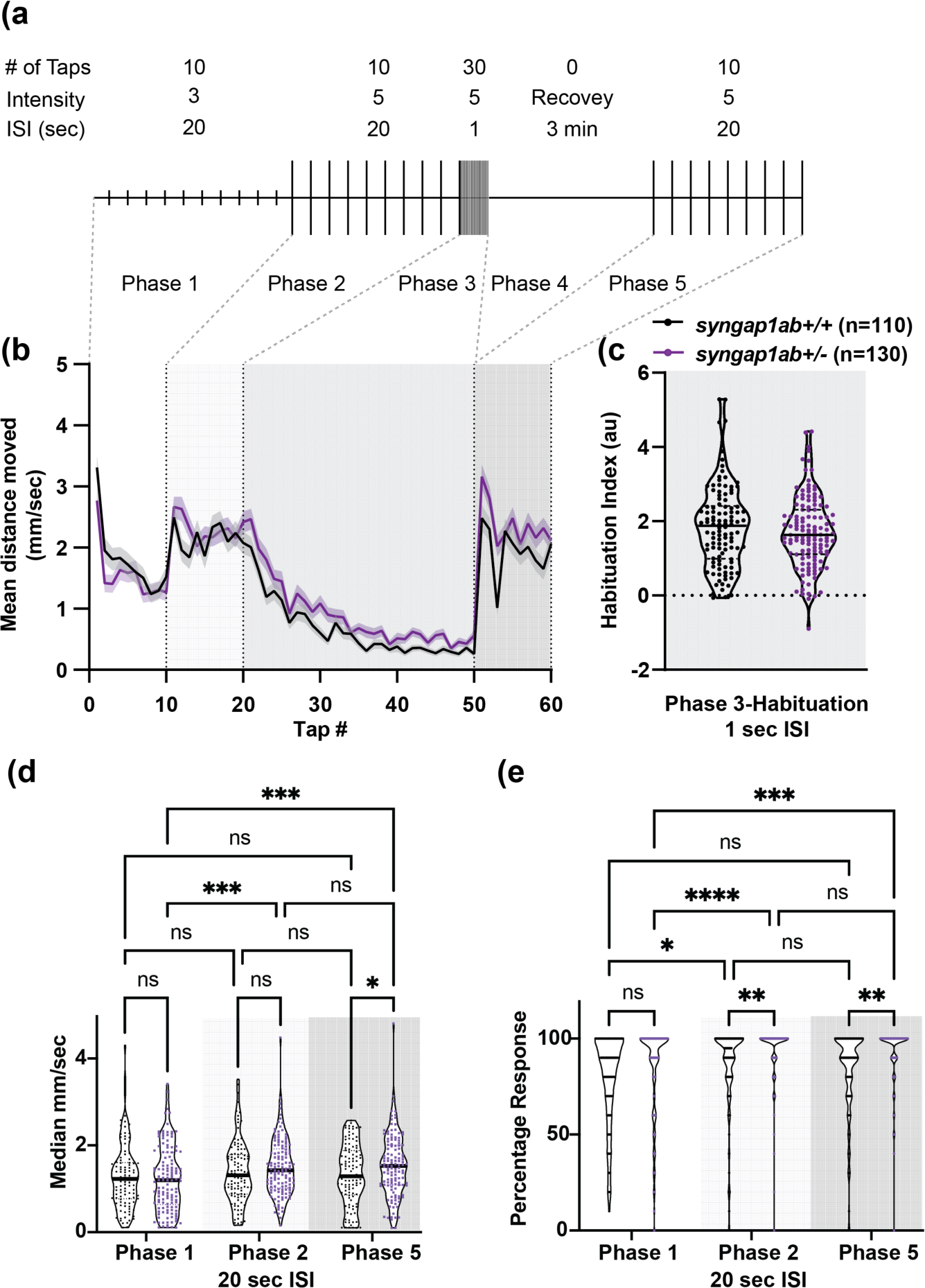
*Syngap1a*b larvae show short-term habituation to vibrational stimuli and more movement in response to stronger stimuli. **a)** A schematic representation of conditions used in the habituation assay (modified from Wolman et al., 2015) **b)** Mean +/- standard error distance moved by WT and *syngap1a*b+/- larvae 1 second post vibration. **c)** Habituation indices was calculated per larva by subtracting the mean distance moved after habituation (taps 41-50) from the mean distance moved pre habituation (taps 11-20). **d)** The Median distance mm/sec moved per phase is shown for WT and *syngap1+/-* larvae. **e)** Percentage response per phase is shown for WT and *syngap1+/-* larvae. A mixed effects model of genotype and phase was conducted and when p<0.05 was followed by a Tukey’s multiple comparison test. * p<0.05; ** p<0.01; *** p<0.001; ****p<0.0001

Despite these similarities, there were differences in that *syngap1+/-* larvae showed consistently elevated responses to high-intensity stimuli during phases 2 and 5. To determine whether this was due to larger movements or an increased probability of response, we calculated median movement per larva (Fig. 3d) and their response probability (Fig. 3e) during Phases 1, 2, and 5. For median movement per second (calculated across taps in a given phase), a mixed effects model of phase and genotype indicated a significant effect of phase (p=0.0018). The subsequent Tukey’s multiple comparison test showing that *syngap1+/-* larvae moved further in response to high-intensity vibrations in phases 2 and 5 than to medium-intensity vibrations in phase 1 (p=0.0006 and 0.0007 respectively) while WT larvae moved similar distances during phases 1, 2, and 5. For probability of response, a mixed effects model indicated effects of both phase (p<0.0001) and genotype (p=0.0009). The subsequent Tukey’s multiple comparison test showed that *syngap1+/-* larvae had a higher response probability to stronger vibrational stimuli (p<0.0001 Phase 1 vs. 2; p=0.0002 Phase 1 vs. 5) and that this higher probability of responses to high-intensity stimuli was greater in *syngap1+/-* than WT larvae (p=0.0049 Phase 2; p=0.0014 Phase 5).

Therefore, in response to stronger stimuli*, syngap1*+/- are more likely to move and move faster, indicating a larger dynamic range of response in *syngap1+/-* larvae. This higher probability of response to the same stimulus is also consistent with higher levels of arousal in *syngap1+/-* larvae.

### syngap1ab mutant hyperactivity is most pronounced in low-arousal settings

We next assessed visual-motor responses (VMR;(Burgess and Granato, 2007; Emran et al., 2008)) in the *syngap1ab* models. For this assay, larval movements are recorded during four cycles of lights-on to lights-off transitions. Larval activity was measured as the total distance moved every 30 seconds. Larvae show robust increases in locomotor activity when presented with a sudden transition from light to darkness (Fig 4a). Compared to WT larvae, both *syngap1ab+/-* and *syngap1ab-/-* larvae showed increased activity (p<0.0001) during both lights-on and lights-off cycles (Fig.4b). During lights-off periods *syngap1ab+/-* showed the greatest movement but this trend was somewhat variable across different batches of larvae (Supplementary Figure 3b); during lights-on, hyperactivity was consistent across batches of larvae and was dependent on the number of mutant *syngap1* alleles with *syngap1ab-/-* showing the greatest movement (Fig. 4; Supplementary Figure 3).

**Figure 4.**
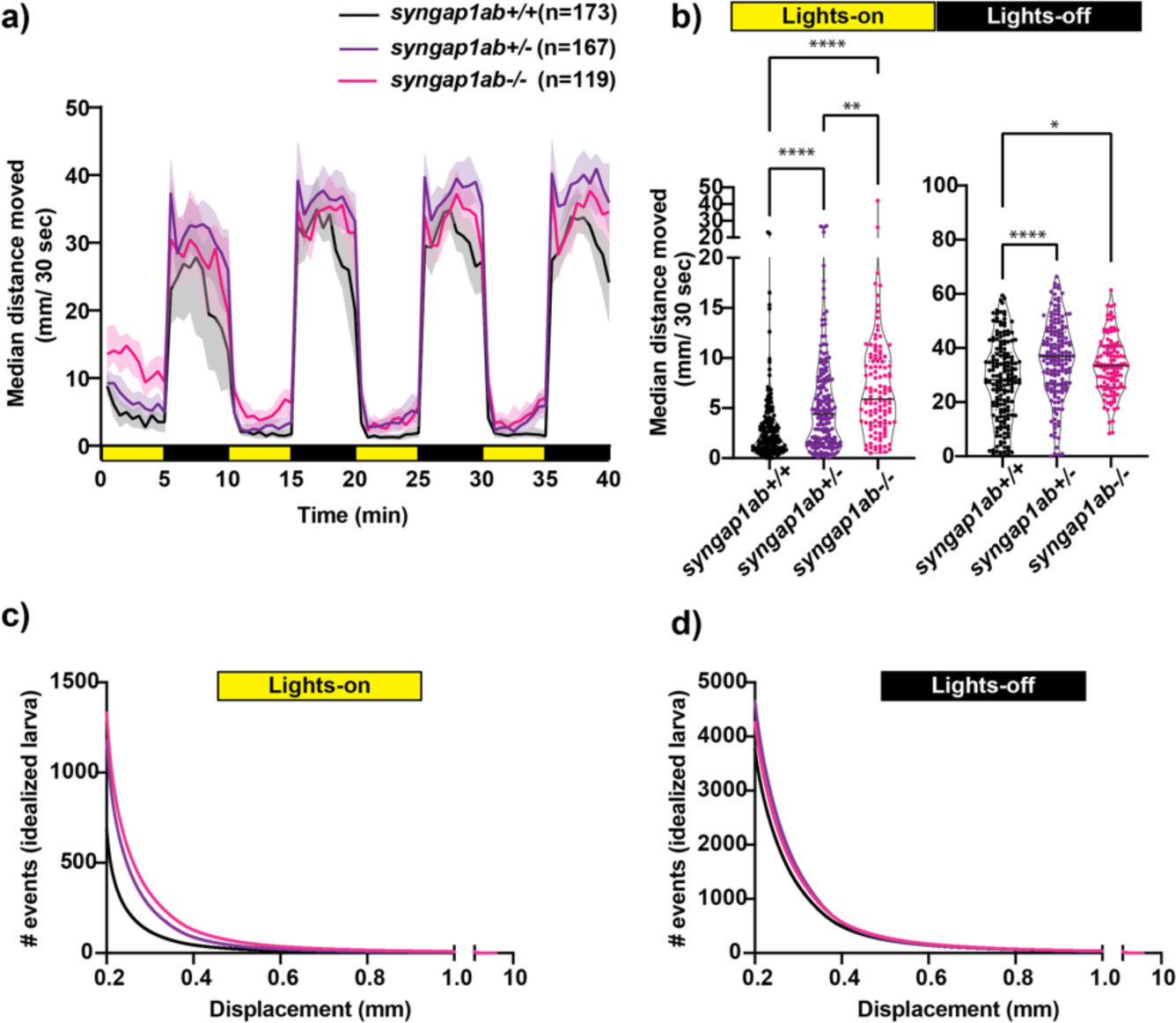
*syngap1* model hyperactivity is most pronounced during light cycles due to a higher frequency of larger movements. **a)** Median± 95% confidence interval distance moved by each 6 dpf larva per 30 seconds, when exposed to 5 minutes of lights-on and 5 minutes of lights-off alternating cycles across five different independent trials (Supplementary Figure 2). **b)** During lights-on cycles, *syngap1ab* mutants showed increased activity levels in a genotype dependent manner where *syngap1ab-/-* were more active than *syngap1ab+/-* which were more active than the WT larvae. During lights-off cycles, syngap1ab mutant larvae showed significantly increased activity compared to WT larvae but there were no significant differences in the activity levels between *syngap1ab-/-* and *syngap1ab+/-* larvae. Statistical analyses between genotypes were carried out using Kruskal-Wallis test followed by Dunn’s multiple comparison test. P value asterisks represent; p<0.05 - *, p<0.01 - **, p<0.001 - ***, p<0.0001-****. Displacement distribution of “idealized larvae” during lights-on **c)** and lights-off **d)** conditions. Graphs were generated by pooling all displacement events during lights-on WT n=119,135, *syngap1ab+/-* n=196,594, and *syngap1ab-/-* n=158,443 and during lights-off WT n=652,368, *syngap1ab+/-* n=777,525, and *syngap1ab-/-* n=507,720 generated from 173 WT, 167 *syngap1ab+/-,* and 119 *syngap1ab-/-* larvae.

For the preceding assays, WT, *syngap1ab+/-*, and *syngap1ab-/-* larvae resulted from independent crosses. To rule out the influence of distinct parental genetic backgrounds on the observed behavior, we examined VMR in larvae resulting from an in-cross between *syngap1ab+/-* adults (Supplementary Fig. 2c-d). Consistent with the previous results, during both lights-on (Supplementary Fig. 2c) and lights-off cycles (Supplementary Fig. 2d), *syngap1ab-/-* had the highest activity levels among all the resulting genotypes.

### syngap1ab hyperactivity in the light resembles WT behavior in the dark

The hyperactivity we observed in the *syngap1ab* mutant models could be due to either increased movement frequency, increased distance traveled per movement, or both. To distinguish among these possibilities, we analyzed the larval movement at a higher temporal resolution. Data were sampled every 25 ms, a much higher temporal resolution than the distance per 30 seconds shown in the preceding figures. Zebrafish movement bouts last ∼250 milliseconds and so 40 Hz resolution is sufficient to capture the majority of bouts with multiple datapoints. We focused on two parameters: the time interval between two consecutive movement bouts, denoted as “dwell time”, and the distance traveled per bout, denoted as “displacement”. These high-resolution activity data were organized and analyzed using custom-written MATLAB scripts to assess the probability of different behaviors, described by displacement and dwell time, during light and dark conditions.

Given highly stochastic individual larval movements, to assess overall patterns of both high and low frequency events across the distribution, we captured a large number of data points from over 100 individuals per genotype and then divided the total bouts by the number of individuals to generate and “idealized larva” for each genotype (Fig. 4b-c). When bouts were pooled across individuals of a genotype, there were >10^5^ bouts per genotype and light condition (Lights-On: WT n=119,135, *syngap1ab+/-* n=196,594, and *syngap1ab-/-* n=158,443; Lights-Off: WT n=652,368, syngap1ab+/- n=777,525, and syngap1ab-/- n=507,720) coming from 173 WT, 167 syngap1ab+/-, and 119 syngap1ab-/- individual larvae. This analysis shows that the differences between genotypes are much more pronounced in the light, low arousal settings, than in the dark, higher arousal setting.

We next looked at probability distributions of dwell times (the time between movements) and distance traveled per movement (Fig. 5). WT larvae in the dark had shorter dwell times and larger movements than WT in the light (Fig. 5ai &ii). These differences in WT light and dark behaviors are highlighted by plots below that show the relative probabilities in Dark versus Light for both dwell time and displacements (Fig. 5aiii &iv). Next, we compared *syngap1ab+/-* and WT in the dark (Fig. 5b) and in the light (Fig. 5c). In the dark, *syngap1ab+/-* and *syngap1ab-/-* had dwell time and displacement distributions that were very similar to WT larvae (Fig 5bi-iv). By contrast in light, *syngap1ab-/-* and *syngap1ab+/-* mutants displayed both more frequent, and larger displacements compared to WT larvae (Fig 5ci & ii). These differences between *syngap1+/-* and WT larvae in the light are highlighted by plots below that show the relative probabilities of *syngap1+/-* (purple) and *syngap1-/-* (pink) versus WT (Fig. 5ciii &iv). These analyses showed that *syngap1*-WT (in light) resembles dark-light comparisons, indicating that *syngap1* mutants behavior in the light resembles that of WT behavior in the dark. Taken together, behavioral experiments show context-dependent hyperactivity that is most subtle during aversive, acoustic stimuli and most pronounced in normally low arousal environments.

**Figure 5:**
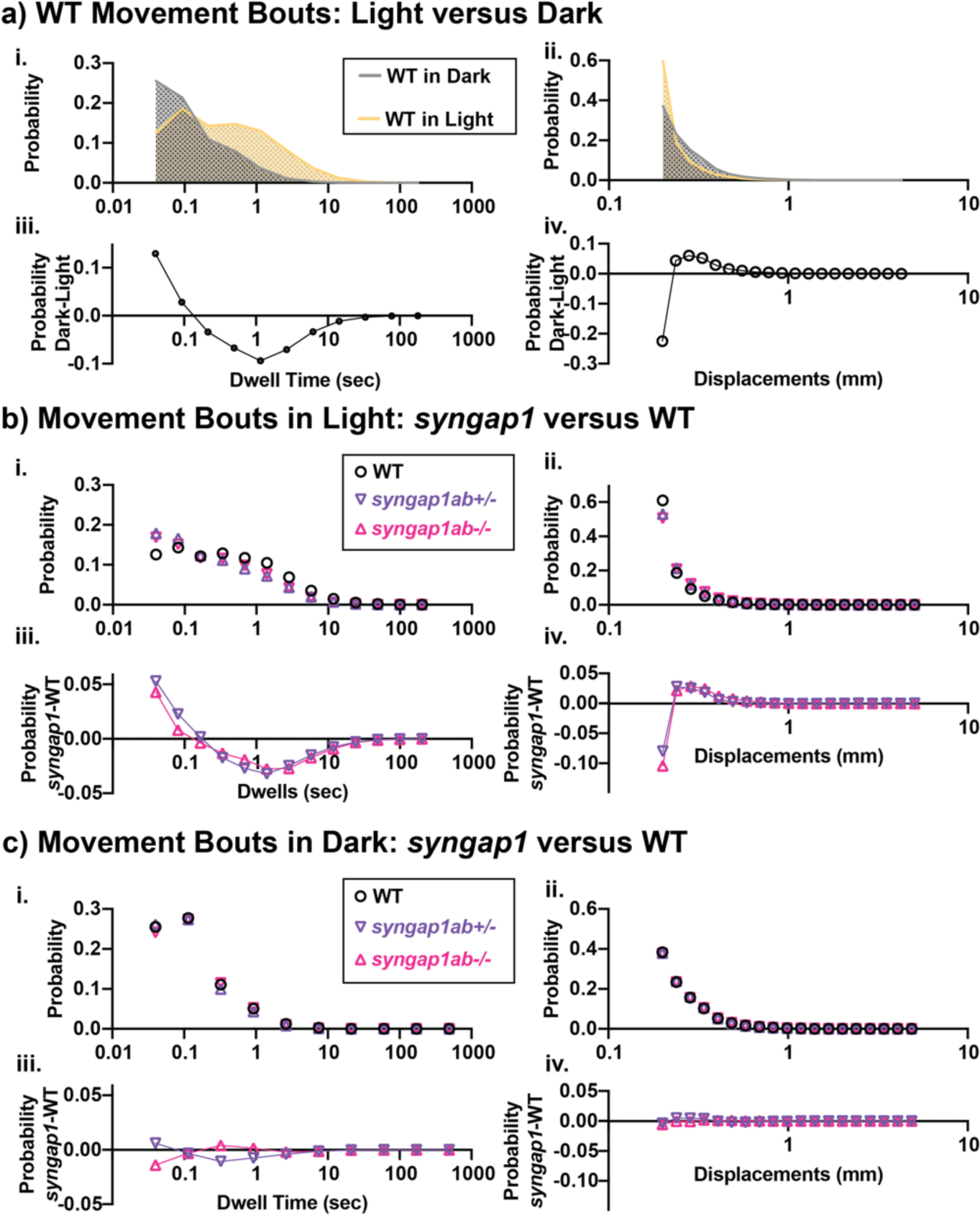
Mutant *syngap1ab* larvae showed heightened arousal during lights-on cycles with more frequent and larger displacements. **(ai & ii)** Probability distributions of dwell times **(ai)** and displacements **(aii)** are plotted for all 173 WT(*syngap1ab+/+*) larvae during lights-on (yellow) and lights-off (dark; checkered) cycles. Below **(aiii & iv)** compare movements in dark and light by plotting probability in dark minus the probability in light. WT larvae moved farther more frequently in dark than in light. **b & c)** Probability distributions of dwell time **(bi & ci)** and displacements **(bii & cii)** are plotted for all 173 WT, 167 *syngap1ab*+/- and 119 *syngap1ab*+/- mutant larvae. Below probability distribution plots **(biii-iv & ciii-iv)** compare movements in *syngap1ab+/-* (purple) and *syngap1ab-/-* (pink) to WT (black) by plotting probability in *syngap1ab* mutants minus the probability in WT. In dark, *syngpa1ab+/-* larvae moved more frequently than either WT or *syngap1ab-/-* larvae while all genotypes had similar displacement distributions. By contrast, in light, both *syngpa1ab+/-* and *syngap1ab-/-* larvae moved more frequently and farther than WT following a similar pattern that WT larvae showed during dark periods, consistent with *syngap1ab* mutants exhibiting a heightened state of arousal in the light. p values were calculated using two-sample Kolmogorov-Smirnov statistical test and for lights on displacement: p(WT vs *syngap1ab+/-*) = 0, p(WT vs *syngap1ab-/-*)=0, and p(*syngap1ab+/-* vs *syngap1ab-/-*)=p<10^-70^) and during lights-off displacement; p(WT vs *syngap1ab+/-*) = 10^-26^, p(WT vs *syngap1ab-/-*)=1.5x10^-11^, and p(*syngap1ab+/-* vs *syngap1ab-/-*)=p<10^-45^), lights-on dwell time (WT vs *syngap1ab+/-*) = 0, p(WT vs *syngap1ab-/-*)=0, and p(*syngap1ab+/-* vs *syngap1ab-/-*)=p<10^-31^) and lights-off dwell time; p(WT vs *syngap1ab+/-*) = 10^-200^, p(WT vs *syngap1ab-/-*)=10^-50^, and p(*syngap1ab+/-* vs *syngap1ab-/-*)=p<10^-200^).

## Discussion

In this study, we generated a zebrafish model of *SYNGAP1* haploinsufficiency and characterized zebrafish *syngap1ab* splice-variants as they relate to mammalian *Syngap1* and zebrafish *syngap1a* and *syngap1b* duplicates. This provided context for the stable zebrafish mutant model of SYNGAP1 syndrome we generated using CRISPR/cas9 genome editing of both *syngap1a* and *synagp1b* duplicates; mutations were validated as loss-of-function alleles at both mRNA and protein levels. We provide evidence that *syngap1a* and *syngap1b* play complementary functional roles in zebrafish with higher larval mortality when *syngap1b* mutant alleles outnumber those from *syngap1a*. Focusing on balanced *syngap1a* and *syngap1b* mutant genotypes for more detailed analyses, we showed that, as in mammalian models and humans, the zebrafish *syngap1ab* models show context-dependent hyperactivity that is especially pronounced in low arousal settings.

Similar to mammalian Syngap1 isoforms (Gou et al., 2020; Yang et al., 2023), both Syngap1 zebrafish orthologs show extensive splicing at both N- and C-termini as well as splice-variants in the middle of the gene that only change a few amino acids. We provide evidence that *syngap1b* but not the *syngap1a* encodes an α1-like isoform, the most extensively studied of the mammalian Syngap1 isoforms (Chen et al., 1998; Kim et al., 1998; Walkup et al., 2016; Araki et al., 2020). In mammals, the α1 isoform has been shown to be enriched at the post-synaptic density of glutamatergic synapses by interacting with the PDZ domain of PSD-95 (Chen et al., 1998; Kim et al., 1998; Komiyama et al., 2002). In mice, loss of the α1 isoform is sufficient to produce cognitive deficits and seizures (Kilinc et al., 2022). The unique expression of the Syngap1 α1 isoform by the *syngap1b* gene may help to explain higher mortality in zebrafish larvae with more *syngap1b* than *syngap1a* mutant alleles. We were not able to determine which of the zebrafish splice-variants corresponded to the other well-characterized mammalian Syngap1 α2, β, and γ isoforms (Kilinc et al., 2018; Araki et al., 2020; Gou et al., 2020), due to sequence divergence in the zebrafish consistent with differences that have been previously described in the zebrafish synapse proteome (Bayes et al., 2017).

Other zebrafish models of SYNGAP1 syndrome mutated only the *syngap1b* gene (Griffin et al., 2021; Colón-Rodríguez et al., 2019). Our differential survival results would predict that the phenotypes reported in *syngap1b* models may relate to a functional imbalance between *syngap1a* and *b*. Our more in-depth subsequent analyses, therefore, focused on larvae with balanced heterozygous mutations in both *syngap1a* and *syngap1b* in an effort to recapitulate mammalian haploinsufficiency.

In humans, altered sensory processing encompasses sensory hyperactivity/ hypoactivity and sensory seeking, and is a core symptom of an ASD diagnosis (Marco et al., 2011; Kirby et al., 2017; Robertson and Baron-Cohen, 2017; Damiano-Goodwin et al., 2018). In SYNGAP1 syndrome specifically, sensory-seeking behaviors include an affinity for contact with flowing water and/or perpetual motion (Wright et al., 2022). Syngap1 haploinsufficiency in both mice and humans alters sensory responses in a way that is context dependent, impacting simple sensory responses, entrainment, and habituation (Carreno-Munoz et al., 2022) and causing increased risk-taking behaviors (Kilinc et al., 2018; Weldon et al., 2018). Further studies on sleep in people with SYNGAP1 syndrome (Smith-Hicks et al., 2021) and mouse models, show disrupted sleep and seizures that are more common at night and often come in clusters that predict transitions between REM and non REM sleep (Sullivan et al., 2020). Taken together, this collection of symptoms suggest a heightened state of arousal and difficulties with behavioral state transitions in people with SYNGAP1 syndrome.

Similar to mammalian models, our *syngap1a*b zebrafish mutant models exhibit context-dependent hyperactivity. In high arousal contexts generated by strong, aversive acoustic stimuli, WT and *syngap1+/-* larvae produced similar highly-stereotyped, high-velocity escape responses, related to startle responses in mammals by their short latency from the stimulus and their dependence on reticulospinal neurons (Liu and Fetcho, 1999; Eaton et al., 2001; Korn and Faber, 2005). In contrast to WT, *syngap1ab+/-* larvae had a larger dynamic range, increasing both distance traveled and response probability with the higher intensity acoustic stimuli compared to WT that had similar responses at both intensities tested. Larger displacements in response to acoustic stimuli have been described in zebrafish glucocorticoid receptor mutants that have chronically elevated glucocorticoids due to a lack of feedback inhibition (Griffiths et al., 2012). It is unlikely that *syngap1ab+/-* mutants have chronically elevated glucocorticoids because elevated glucocorticoids reduce movement in the light, unlike our *syngap1ab+/-* model that moves faster and more frequently in the light.

Another possibility is that other arousal pathways, groups of neurons and/or neurotransmitters known to promote behaviors, are more active in the *syngap1* model. In both mammals and zebrafish, arousal pathways including dopamine, QRFP, serotonin, and hypocretin/orexin among others have been linked to increased locomotion (Chiu and Prober, 2013; Lovett-Barron et al., 2017; Corradi and Filosa, 2021; Tan et al., 2022). Moreover, gain-of-function experiments in zebrafish have shown that overexpressing either hypocretin/orexin or CART (cocaine and amphetamine regulated transcript) is sufficient to increase the probability that zebrafish larvae will respond to acoustic stimuli (Prober et al., 2006; Woods et al., 2014); and overexpression of any one of hypocretin/orexin, calcitonin gene related peptide (cgrp), or cholecystokinin (cck) is sufficient to increase daytime movement frequencies (Woods et al., 2014). Thus, overactivation of select arousal pathways is consistent with *syngap1ab+/-* behavioral profiles.

Tests of behavioral responses to alternating light and dark conditions showed that *syngap1+/-* hyperactivity was greatest in the light, a setting characterized by long dwell times and short movements in WT larvae. In contrast to WT, *syngap1ab+/-* and *syngap1ab-/-* larvae had more short dwell times and larger movements, patterns much like those seen in WT when they are suddenly transitioned to the dark as in the VMR assay (Burgess and Granato, 2007; Emran et al., 2008; Kozol et al., 2021). Increased WT movements with sudden darkness have been shown to reflect goal-directed behaviors as the fish seek to return to the light, a behavior known as dark phototaxis (Horstick et al., 2017). Goal-directed increases in activity can also result from internal states, such as hunger, studied in 7-day-old zebrafish larvae that no longer have a yolk supply (Filosa et al., 2016; Wee et al., 2019). In this study, hyperactive *syngap1ab* mutants were 6-day-olds and therefore their hyperactivity was not likely to be driven by hunger. Hunger-induced hyperactivity is associated with reduced cortisol, increased activity in the serotonergic raphe neurons and increased risk-taking as larvae approach objects that could be either food items or predators (Filosa et al., 2016). Like hunger, flowing water can evoke more frequent movements that are also dependent upon dorsal raphe serotonergic pathways (Yokogawa et al., 2012). It is possible that the hyperactivity that we observe in the *syngap1ab+/-* larvae is a form of sensory seeking behavior, reminiscent of perpetual motion and affinity for flowing water seen in people with *SYNGAP1* syndrome.

Studies in people with genetic forms of autism and in animal models show that, despite some shared aspects of etiology, including changes to neuro- and gliogenesis and altered excitatory/inhibitory balance (Hoffman et al., 2016; Willsey et al., 2021), detailed changes in neuroanatomy, behavioral profiles, and associated symptoms exhibit substantial differences by genotype supporting the existence of autism subtypes (Ellegood et al., 2015; Thyme et al., 2019; Zerbi et al., 2021; Weinschutz Mendes et al., 2023). In the case of *SYNGAP1*, both human and animal model studies point to arousal pathways as playing an important role in context-dependent hyperactivity.

## Supporting information

Data used for Figures

Data Analyses

## Acknowledgements

We would like to thank the SYNGAP1 Foundation, the SYNGAP1 Research Fund, and individuals with SRID and their families for their support and advocacy. We thank Millie Rogers, Adhikansh Jain, and Dr. James Baker for their valuable feedback that improved the manuscript. We would like to acknowledge the excellent care of the zebrafish provided by the University of Miami Zebrafish Facility manager Ricardo Cepeda. This work was supported by Bridge Funds from the College of Arts and Sciences at the University of Miami and NIH grants NIMH R03MH103857, NICHD R21HD093021, and SFARI Pilot grant 719401 to JED, NIH grants MH112151 and NS03671 to RLH, and NSF IOS#2131037 to SS.

---

**Supplementary Figure 1.**
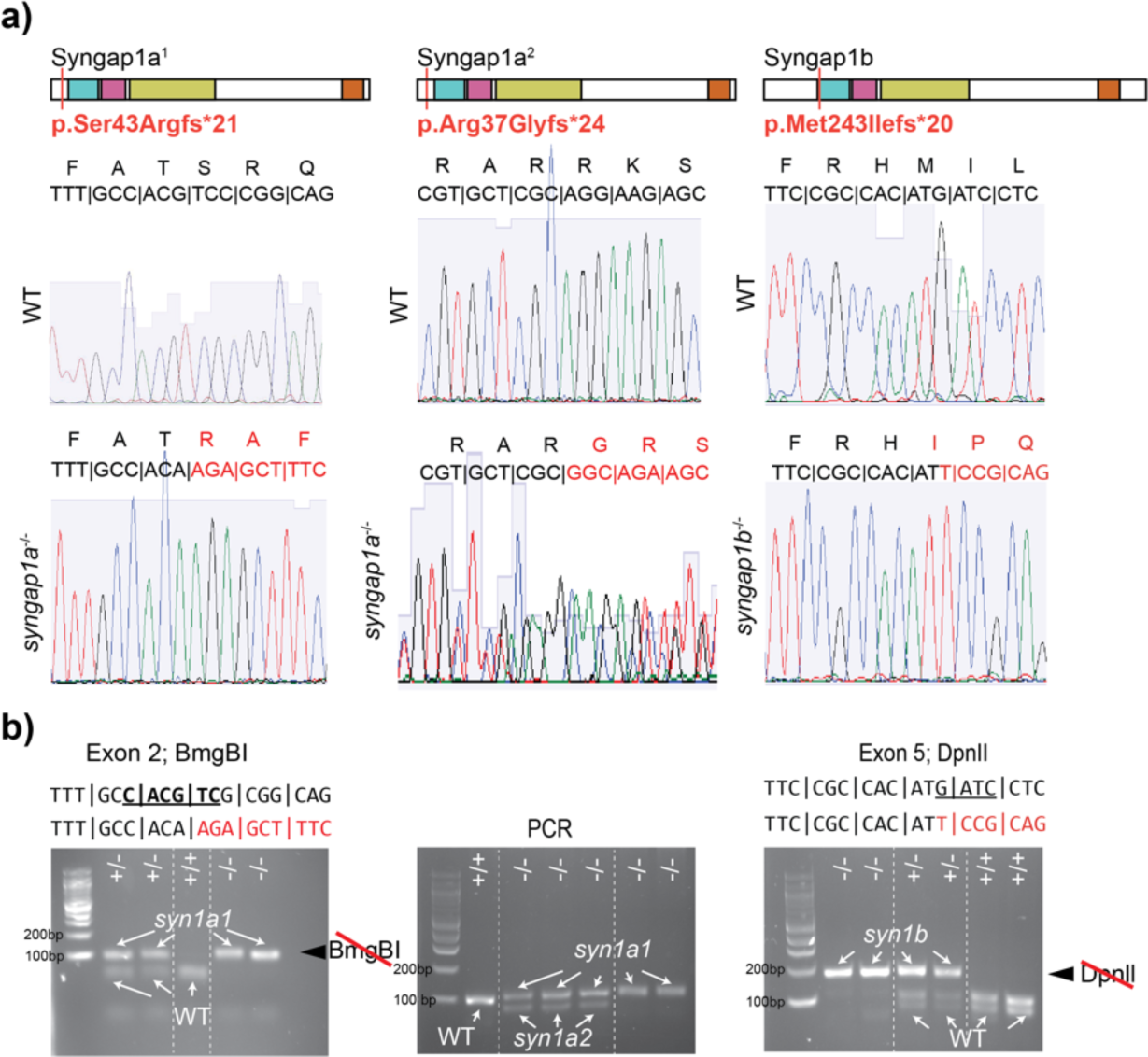
Genotyping zebrafish *syngap1a* and *syngap1b* mutant alleles. **a)** Syngap1a and b protein diagrams are shown, one for each mutant allele, with red vertical lines indicating the location of each frame-shift mutation. Red type below the lines provides the details of the frameshift indicating the position of original amino acid, what it was changed to and the subsequent number new amino acids before the stop codon truncation. Below each diagram are two electropherograms, the top showing the WT and the bottom showing the mutant sequences. Codons and corresponding amino acids shown above each electropherogram. **b)** Genotyping gels are shown for each allele. For *syngap1a1*, a stretch of exon 2 that includes a restriction enzyme site for BmgB1 in the WT sequence is destroyed in *syngap1a1*. *Syngap1a2* has a 22 base deletion that can be detected as a size shift directly from PCR amplification of Exon 2 surrounding the mutated site. For *syngap1b*, a stretch of exon 5 that includes a restriction site for DpnII in WT is destroyed in the *syngap1b* mutant allele.

**Supplementary Figure 2.**
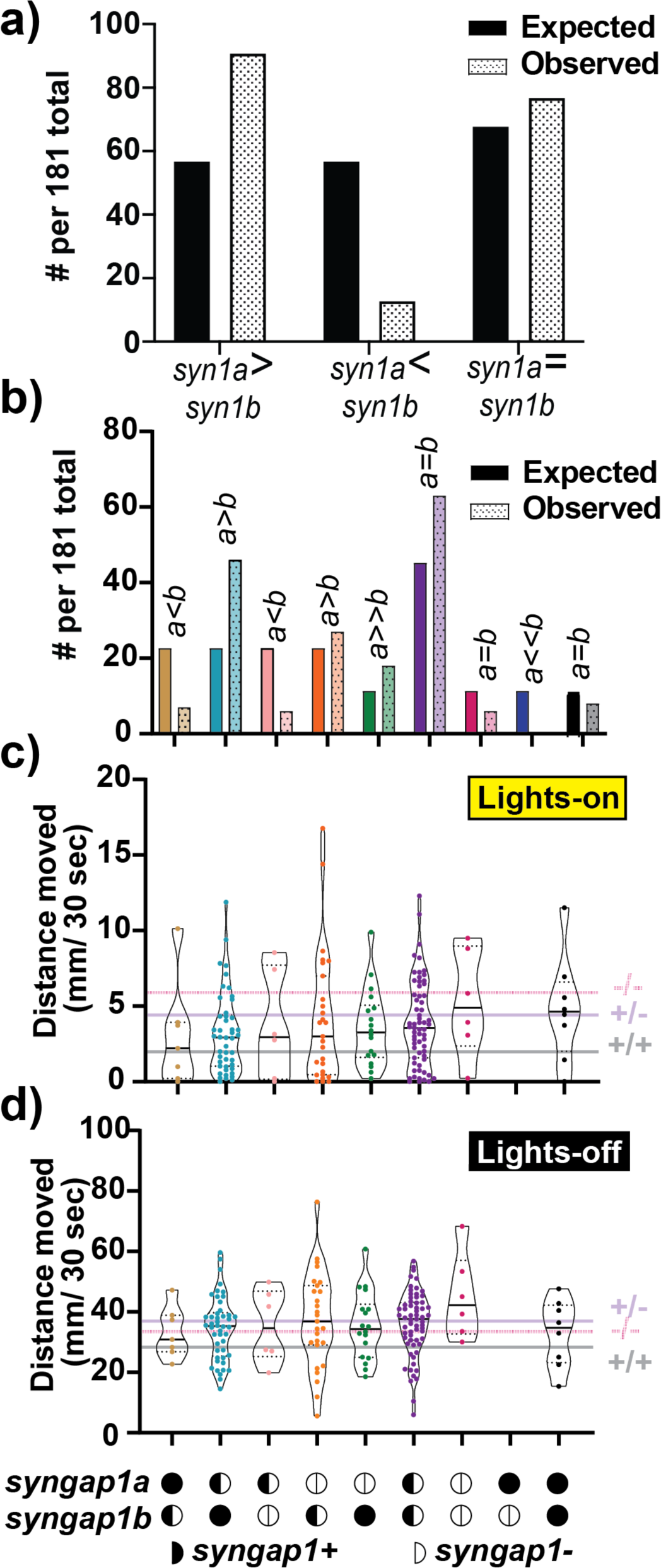
*Syngap1b* is more important for larval survival than *Syngap1a.* **a & b)** Expected vs observed ratios of different *syngap1a* and *syngap1b* mutant allele combinations resulting from a *syngap1a*b+/- in-cross. A Chi-square test indicates that observed is different from expected ratios p<0.0001. **a & b)** Of note, allele combinations for which *syngap1b* mutant alleles outnumber those of *syngap1a* (a<b) are under-represented with the most extreme case being no surviving larvae with the *syngap1a+/+; syngap1b-/-* genotype. Median distance traveled in the light **c)** and in the dark **d)** are plotted against genotype for all allele combinations with the highest median distances seen in *syngap1a-/- ;syngap1b-/-* larvae. Horizontal lines indicate medians from other analyses to contextualize values. Because of low numbers in some genotypes, these medians may not be representative.

**Supplementary Figure 3.**
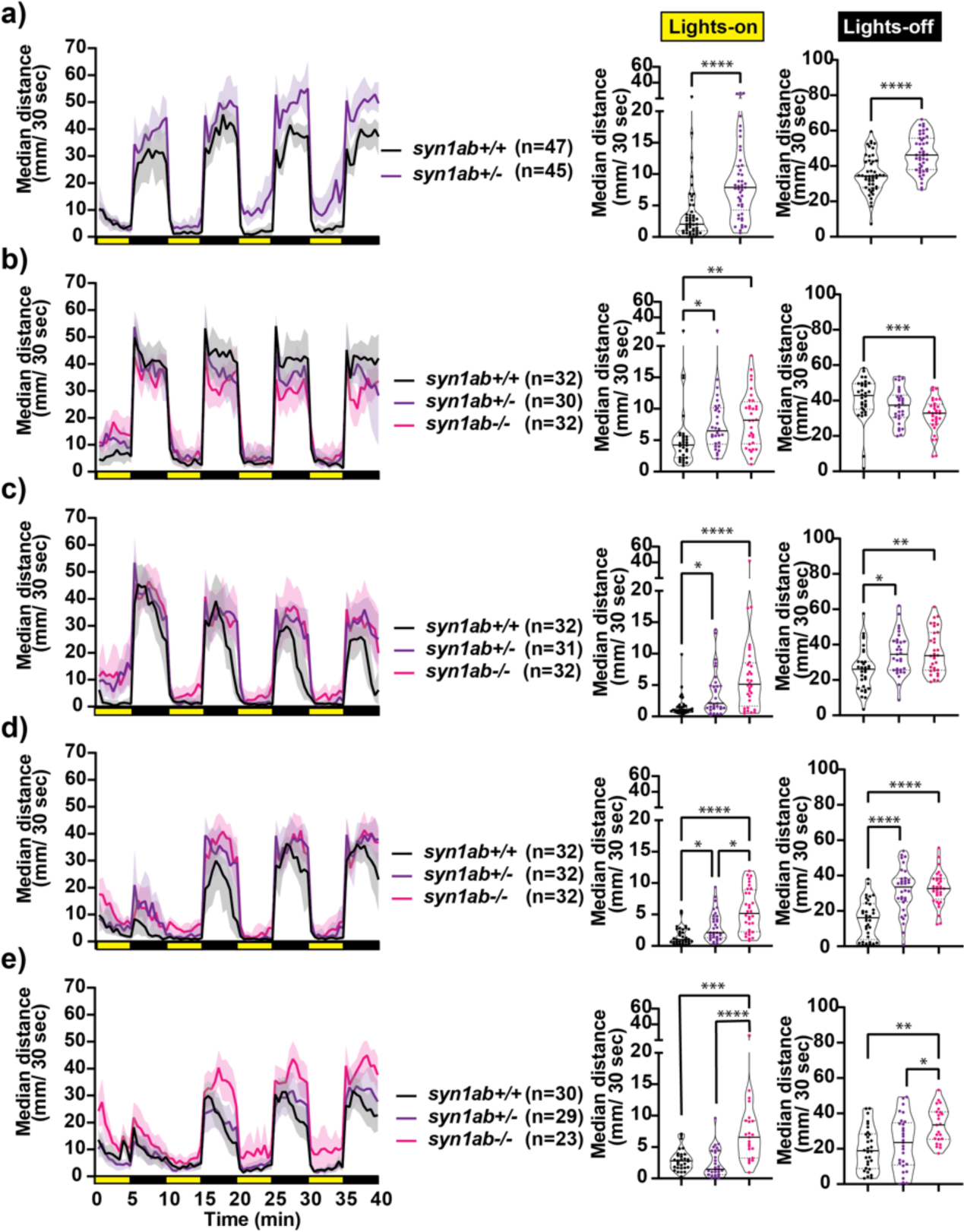
Visual Motor Response data per batch shows *syngap1* is more consistently hyperactive in the light than in the dark. **a-e)** Median ± 95% confidence interval distance moved by each 6 dpf larva per 30 seconds, when exposed to 5 minutes of lights-on and 5 minutes of lights-off alternating cycles across five different independent trials sample size is indicated to the right of each VMR plot. To the right of that genotypes are compared in light and dark separately. In lights-on cycles, *syngap1ab* mutants showed increased activity levels in a genotype dependent manner where *syngap1ab-/-* were more active than *syngap1ab+/-* which were more active than the WT larvae. During lights-off cycles, syngap1ab mutant larvae often but not always showed significantly increased activity compared to WT larvae (see b). Statistical analyses between genotypes were carried out using Kruskal-Wallis test followed by Dunn’s multiple comparison test. P value asterisks represent; p<0.05 - *, p<0.01 - **, p<0.001 - ***, p<0.0001-****.

